# A Network Approach to DNA Methylation Clocks

**DOI:** 10.64898/2026.06.18.733218

**Authors:** Antón Carcedo, Seong-Gyu Yang, Jelena Smiljanić, Magnus Neuman, Sigrid Wennstedt, Sofie Degerman, Ludvig Lizana

## Abstract

Biological age predicts health and lifespan better than chronological age, but remains difficult to measure. One leading molecular proxy for biological age is DNA methylation, which underlies age predictors known as “clocks”. These clocks use penalized linear regression to predict chronological age from methylation levels using selected cytosine–guanine pairs (CpGs) along DNA. Although they predict chronological age within a few years and track mortality risk, there are several issues. Different clocks share a vanishingly small number of CpG sites, many of which show weak associations with age. Also, the clocks often do not transfer across methylation array platforms. This paper takes a network approach to better understand these issues. By using 12 public datasets from human blood, we build a co-methylation network of the sites that show the strongest age correlation. After pruning weak links, we find that it has a small number of large modules of covarying CpGs surrounded by many small modules and singleton sites. These modules are biologically interpretable, as they are associated with CpG island contexts and enriched for distinct Gene Ontology functions. We also map five established clocks onto this network (Horvath, Hannum, AltumAge, Skin & Blood, and Han) and find that they select some CpGs from the same module. This suggests that they are more similar than they appear. The network structure also suggests new ways to build clocks. A simple clock that retains one CpG per module matches the performance of established clocks. A second one, built from module-level principal components, outperforms all five established clocks in three validation cohorts and is transferable across array platforms (Illumina Infinium Methylation 450K or EPIC arrays). Overall, the network perspective shifts attention from individual CpG sites to modules of covarying sites. This perspective helps explain why DNA methylation clocks perform so well despite their differences and provides a more systematic approach for developing the next generation of aging biomarkers.

## 1. INTRODUCTION

While chronological age simply records time elapsed since birth, biological age reflects one’s physiological state^1^, such as wear and damage that accumulate across tissues, cells, and molecules during aging^2^. Its pace also varies substantially across individuals^3^. Because aging arises from many interacting processes rather than a single cause, biological age is difficult to measure directly. Researchers have therefore developed biomarkers that aim to integrate multiple aging processes into a single quantitative measure. Among the proposed biomarkers, DNA methylation (DNAm) has emerged as one of the most promising candidates, leading to the development of epigenetic aging clocks^4^.

Most DNAm clocks are made from penalized linear regression models that use DNAm array data covering 450,000 to 850,000 CpG probes distributed across the genome^5–7^. After extensive data filtering, researchers have selected a subset of CpG sites and computed a weighted sum of DNAm values to construct DNAm clocks. Estimating biological age using these clocks shows strong correlations with chronological age, health, and mortality risk^8;9^. As a result, researchers increasingly use them to study accelerated aging and its associations with genetic, environmental, and lifestyle factors. DNAm clocks have also found applications in less conventional settings. For example, forensic scientists use them to estimate the age of potential perpetrators from crime-scene bio-samples^10^, and immigration authorities apply DNAm clocks to estimate the age of refugees lacking valid identity documents^11^.

However, current DNAm clocks also have limitations^12^. As mentioned before, most clocks rely on linear regression methods (e.g., Lasso, Ridge, or Elastic Net) or Neural Networks to select CpG sites from high-dimensional methylation data. Although these approaches produce accurate predictors, they often identify CpG sets that are difficult to interpret biologically. In addition, clock CpG sites do not translate well across methylation array platforms, because some probes are no longer covered^13^. Different clocks also share surprisingly little overlap in their CpG composition, despite having comparable predictive performance. As a result, it remains unclear why DNAm clocks work so well, and why many clock-associated CpG sites lack clear biological interpretation.

To address some of these limitations, we develop a network approach to DNAm clocks. The method constructs a co-methylation network that groups CpG sites into modules with similar methylation variation across a sample group with several individuals. By focusing on these modules, the approach captures relationships among CpG sites that existing clocks ignore, as these clocks typically treat the sites as independent elements. Our framework has two main advantages. First, it allows researchers to analyze clock-associated CpG sites within a network topology alongside common bioinformatic features such as chromatin states and gene annotations. Second, it provides a more systematic way to select CpG sites to build new age predictors without relying on opaque machine-learning methods. We demonstrate this using several DNAm datasets from blood and construct a network clock that achieves higher accuracy than any other established DNAm clock.

## II. METHODS

### DNA methylation data sets

DNAm can be measured by a variety of methods, but is commonly measured with BeadChip arrays. These arrays use probes to measure the presence or absence of -groups at specific cytosines located in genomewide distributed CpG sites. For each CpG site, the arrays report light intensities that can be used to estimate its proportion of methylation *β:*

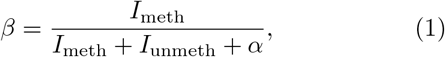

where *I*_meth_ and *I*_unmeth_ are the intensities of the methylation and unmethylation probes, respectively, and *β* is a constant that stabilizes *β* at low signal intensities. We set *α* = 100, consistent with standard Illumina array processing.

We build our networks and clocks from human blood. We use a single tissue because DNA methylation patterns differ between tissues, which keeps the age-related changes we study from being blurred by tissue-to-tissue differences. We choose blood because it is the most widely profiled tissue for DNAm, with many public datasets available. We combine 12 of these datasets to perform our analysis and train our clock rather than relying on a single cohort. Pooling spreads the data across Different populations and batches, so the methylation changes we keep reflect age broadly, not the conditions of any single cohort. The 12 training datasets are GSE42861^14^, GSE51032^15^, GSE59065^16^, GSE61496^17^, GSE81961^18^, GSE87571^19^, GSE87640^20^, GSE87648^20^, GSE99624^21^, GSE107737^22^, GSE125105^23^, and E-MTAB-4931^24^. These cohorts were profiled on Illumina 450K arrays, which report *β*-values at 450,000 CpG sites.

From these datasets we keep only healthy control samples. Because we want to capture the methylation changes that track chronological age, restricting the analysis to healthy individuals avoids the extra variation that disease introduces, so the networks reflect aging rather than a mix of aging and illness. The resulting training dataset contains 1917 samples aged 18 to 94 years, with a 61% female proportion.

We also construct a validation set from three Illumina EPIC arrays (EPIC v1, 850K): GSE200376^25^, GSE217633^26^, and GSE235717^27^. From these cohorts, we include patients as well as healthy controls. This lets us test how the clocks predict age in the presence of disease, here HIV, psoriasis vulgaris, and psoriatic arthritis. For GSE200376 and GSE235717, all samples (including patients) are included, whereas for GSE217633, we include all 44 controls and the same number of randomly selected HIV patients. The validation set contains 187 samples from individuals aged 19–77 years.

## Data preprocessing

To analyze DNAm array data, we first perform signal corrections, probe filtering, and normalization for each sample. These steps reduce the technical biases and make the measurements comparable across samples^28^. We merge the cohorts into a combined dataset so that the patterns we find reflect age across many studies rather than the particularities of any one. Because the cohorts come from Different studies, this merged data carries batch effects, i.e., technical systematic differences. We remove these with ComBat, so the samples can be analyzed together. For the validation set, we follow the same procedure without applying ComBat. Leaving it uncorrected keeps it independent of the training data and mimics how the clock would be used on genuinely new data.

### Signal correction

We implement signal correction using the R package `minfi` ^29^. Raw intensity measurements from Illumina arrays are subject to background noise and dye bias inherent to the two-color fluorescence system. To correct for these artifacts, we apply the normal-exponential out-of-band (Noob) algorithm^30^. The Illumina 450K and 850K arrays have two types of probes: type-I probes use only one of two color channels, whereas type-II probes use both green and red channels. Since the scanner always measures both channels at every probe, the unused channel of a type-I probe captures only background noise. Such noise signal from type-I probe is called the out-of-band signal, and is used to estimate and remove the background noise. Noob also corrects for dye bias between the red and green channels. Assuming that type-I probes from each channel share similar overall intensity distributions, a scaling factor between red and green channels is estimated from these probes and applied across all probes.

### Probe filtering and normalization

After the background noise and dye-bias correction using the Noob algorithm, we filter out probes and normalize the beta values using R package `ChAMP` ^31^. In the probe filtering process, we remove CpG sites on sex chromosomes, probes overlapping known single nucleotide polymorphisms, probes with detection p-value greater than 0.01, and multi-hit probes. After probe filtering, 336,983 CpG sites (∼ 337k CpGs) remain.

Next, we apply the beta-mixture quantile (BMIQ) normalization to the remaining probes^32^. BMIQ is a within-sample method that corrects for probe design bias by adjusting the *β* distribution of type-II probes to match that of type-I probes.

### Batch-effect removal

Before pooling the 12 training datasets, we remove batch effects that would otherwise be mistaken for biological signal. This process is implemented by using R package `sva` ^33^. We apply ComBat^33;34^, which provides a statistically robust framework for batch effects correction while preserving biological covariates of interest. In this study, we do not include biological covariates such as age, and sex as preliminary tests show that including them as covariates produces nearly identical *β* values to the covariate-free results (S1 Fig).

In this step, we first assemble the *β-*values for around 337k CpGs across 1917 samples, after probe filtering and BMIQ normalization. We then transform the *β* values into *M* values as follows:

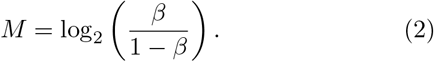

The *M* value exhibits a more symmetric distribution with improved variance stability compared to *β*. Due to computational constraints, we apply ComBat to the *M* -values in chunks of 10k CpGs. After applying ComBat for all chunks, we transform *M* back to *β* to construct our clock. We verify that, after removing batch effects, there are no dataset-specific modules using PCA (S2 Fig).

### Constructing co-methylation networks

We analyze genome-wide DNAm levels by constructing a co-methylation network. In this weighted network, CpG sites are nodes, and the link weight between two sites is the Spearman correlation between their methylation levels. Full methylation arrays produce large and dense networks with no modular structure. For example, the 450K array contains about 450,000 CpG sites and the corresponding network therefore ∼10^11^ links. To reduce the network size, we restrict the analysis to CpG sites associated with age and then prune weak links. Below, we outline the general procedure and the following sections provide detailed explanations.

- **Remove CpGs uncorrelated with age**. For each CpG site *i*, we first compute the Spearman correlation *α*_*i*,age_ between *β*_*i*_ values and chronological age. Next, we retain sites with |*ρ*_*i*,age_| values above a cutoff of 0.35, resulting in a network with 7, 328 nodes. Replacing Spearman with Pearson correlation yields a quantitatively similar result (S3 Fig).
- **Calculate link weights**. For every pair of remaining CpGs, we compute the Spearman correlations between their *β* values. This yields a fully connected network, exemplified in Fig. 1a.
- **Prune links**. The correlation network is dense and has no modular structure. We therefore prune insignificant links using a threshold method (Fig. 1b).
- **Find modules**. To identify modules, or groups, of co-varying CpGs, we cluster the pruned network with the Infomap algorithm^35–38^. Infomap assigns each CpG to a module so that within-module connections are dense and between-module connections sparse (Fig. 1c).

**FIG. 1.**
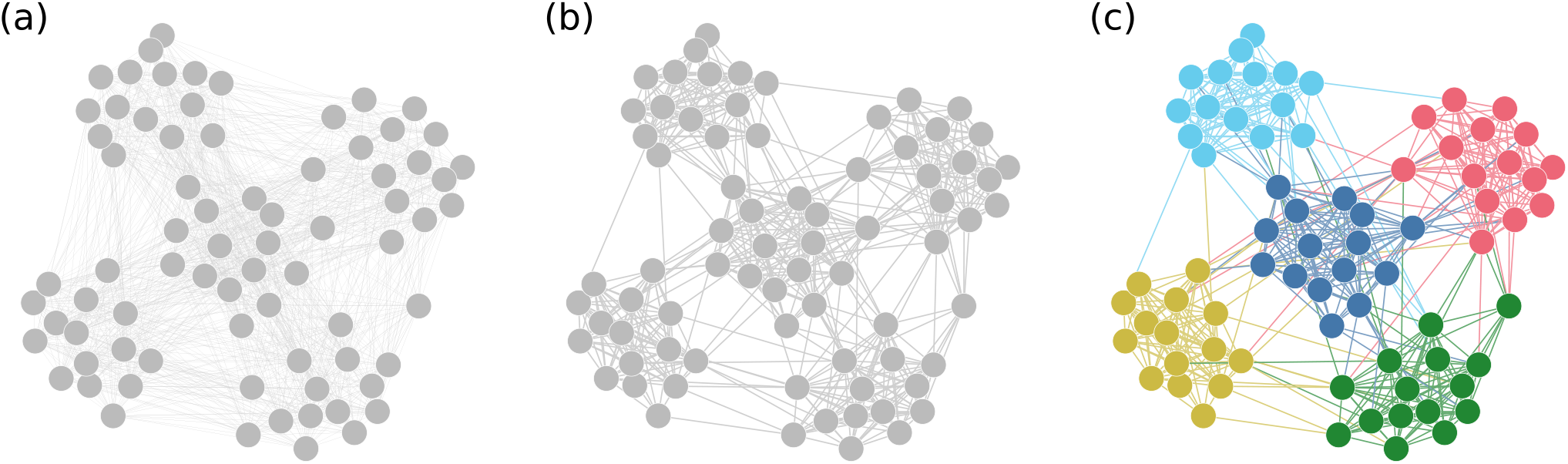
Constructing co-methylation networks from DNAm array data. (a) First, we compute CpG–CpG correlations (Spearman) across samples to form a weighted network (nodes = CpGs; edges = correlation values). (b) Second, we prune links using a thresholding technique to separate signal from noise. (c) Third, we find modules of co-varying CpGs using `Infomap` .

### Network link pruning

By construction, correlation networks are dense because the correlation is never exactly zero. Therefore, we must remove links to reveal the underlying structure. We use thresholding, one of numerous options^39^, where we construct an unweighted, undirected network in which two CpGs (*i* and *j*) are connected only if their correlation exceeds a threshold τ . Mathematically, the new adjacency matrix becomes

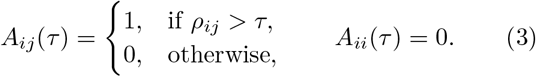

We keep only positive correlations, dropping negative ones which we find to be weaker in our dataset. We do this because we want modules of CpGs that vary together, sites whose methylation rises and falls in lockstep from sample to sample. When CpGs move together this way, any one of them can represent the shared change. Negatively correlated sites do the opposite, moving in opposite directions, so one cannot stand in for the other, and we leave these links out. Next, we explain how we determine the threshold *τ*.

### Threshold selection

A robust threshold *τ* gives the same module composition even under perturbations of the data. To assess robustness, we bootstrap the samples 100 times. For each bootstrap dataset, we construct the network at different thresholds *τ* and infer the network partition with Infomap^35–38^. We then measure how similar partitions are across bootstraps using the adjusted mutual information (AMI). As *τ* increases from 0.3 to 0.9 the network breaks into more modules and leaves more singletons, risking overfitting (S4 Fig). We find that the AMI between bootstraps drops when *τ* ≈ 0.8. We therefore restrict the thresholds to 0.65 *≤ τ* ≤0.75, which is high enough to split the largest module but low enough to avoid overfitting. Our criterion is stricter than standard significance testing, since a correlation this strong lies far beyond the thresholds that standard corrections would require^40^. Within this range, the overall topology stays consistent, where a small number of large modules are surrounded by a sea of singletons (S5 Fig).

Modules corresponding to *τ* = 0.7 recover more and cleaner functional themes than those at adjacent values (S5 Fig). Due to the biological coherence of the resulting modules and stability across bootstraps, we use *τ* = 0.7 in the subsequent analysis.

### Clustering networks with Infomap

To detect modules of co-varying CpGs, we cluster the pruned network using Infomap, a widely used network community detection method that performs well in benchmarks^41;42^. Infomap uses an information-theoretic approach and is particularly suited for correlation networks with high clustering coeffcients^43^. We run standard Infomap, allowing for hierarchical clustering. Nodes left isolated by pruning are assigned to their own singleton module. The final output is a network partition of modules (see Fig. 1c.)

### Gene ontology analysis

To test whether the network modules correspond to distinct biological processes, we run gene ontology (GO) enrichment analysis^44^ on each module separately. We run it in R with the `GOmeth` function from the `missMethyl` package^45^. Built for methylation arrays, GOmeth corrects two biases that would otherwise inflate enrichment, the deffering numbers of probes per gene and CpGs annotated to several genes. For the gene annotations, it draws on `org.Hs.eg.db` ^46^. Each module is then tested against a common background, all 7,328 CpGs in the co-methylation network.

This test returns the GO terms enriched in each module. We keep the terms significant at a nominal level (*p <* 0.05) and group related ones into parent terms with the `rrvgo` package^47^, at a semantic-similarity threshold of 0.7. We then report the parent terms by their nominal enrichment *p*-value. No term survives the stricter correction for multiple testing, which is expected for modules of only tens of CpGs. We therefore treat the enrichment as exploratory and report the parent terms by their nominal p-value, using them to compare functional themes across modules rather than to single out individual terms.

### Building network clocks

Having found that age-associated CpGs organize into modules, we develop a method that uses these modules as building blocks of an age predictor. The modules contain CpGs that co-vary strongly, suggesting that a single representative site should reflect the methylation state for the entire module. To test this, we build a clock that selects *N* CpGs, one from each module. As a reference case, we also randomly select *N* sites from the entire network, thereby allowing oversampling of some modules. We denote these two strategies as:

1. **Random sampling**. Draw *N* CpGs at random from the full age-filtered network.
2. **One-per-module sampling**. Draw *N* CpGs at random under the constraint of at most one CpG per module.

Next, we create an age predictor 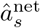 of sample *s* as

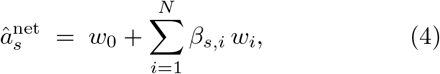

where *β*_*s,i*_ is the *β*-value of CpG *i* in sample *s*, and *w*_0_,. .., *w*_*N*_ are weights.

We calculate the weights with Ridge regression. With up to *N* = 500 CpGs and a limited number of training samples, ordinary least squares tends to overfit, giving coeffcient estimates with high variance that generalize poorly. If selected CpGs are correlated, as can happen under random sampling, this high variance problem is worse. Ridge adds a penalty on the squared coeffcients, which shrinks the weights and trades a little bias for lower variance, giving estimates that generalize better. The penalty also sets ridge apart from the lasso or elastic net often used for established clocks. Those drive some weights exactly to zero, performing CpG selection, whereas ridge keeps all *N* CpGs. Here, the selection is already done by the network, so we want a method that weights the CpGs rather than discards them.

We train and evaluate each clock using 5-fold cross-validation on a pool comprising 12 datasets. We report the average test performance across folds. Repeating the random sampling many times yields an ensemble of clocks, which lets us measure how much performance depends on the particular CpG set. Finally, we test the best clock on the held-out validation set.

### Building a module-level PCA clock

The two network clock strategies above select CpGs at random. While they perform similarly on average, the realizations differ slightly, and we cannot predict which one will perform best on the validation set. To mitigate this variability, we slightly modify the data representation by using a single methylation value for each module, such as the median or average. In practice, we go through each row in the array matrix, i.e, each sample *s*, and group CpG sites that belong to the same module *c*. Next, we calculate the summary statistic for each module in each row, *x*_*s,c*_, and create a new matrix *X* with size (# samples) × (# modules).

While this new data set offers no alternatives for within-module choices, it poses another problem: Which data points should one pick, and how many? To circumvent this issue, we use Principal Component Analysis (PCA) to extract the dimensions that explain most of the variation in the data and to build the clock from the leading components. We define our new age predictor, the “PCA clock”, as

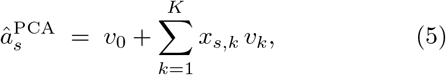

where 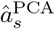 denotes the predicted age, *v*_*k*_ are the weights, and *K* is the number of PCs we choose to include in the sum (typically we omit most). We determine the weights *v*_0_,. .., *v*_*K*_ using Ridge regression and a training dataset. For completeness, prior to calculating the PCs, we followed standard protocol and normalized each column in the *X* matrix to zero mean and unit variance.

Once the weights are determined, it is straightforward to apply the PCA clock to a new dataset. First, assemble the methylation values in that dataset for the CpGs associated with each module and compute the medians (S6 Fig). Next, use these as *x*_*s,c*_ in the sum in Eq. 5 and calculate 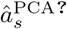 .

## III. RESULTS

### Limitations of existing DNAm clocks

Established DNAm clocks share very few CpG sites. Yet, even if using Different parts of the methylome, they predict age accurately. To illustrate the differences, we analyze five widely used chronological-age clocks: AltumAge^48^, Hannum^5^, Horvath^6^, Skin & Blood^49^, and Han^50^ (Fig. 2). To quantify the overlap between them, we compute the Jaccard index *J* (the number of CpGs shared by two clocks divided by the size of their union). The overlap is small between most clocks(*J* ✢ 0.11; Fig. 2a). An exception is Hannum and Han (*J* = 0.15), which likely reflects that they both were trained on blood data. With such a small overlap, how can clocks built from largely disjoint CpG sets all predict age so well?

**FIG. 2.**
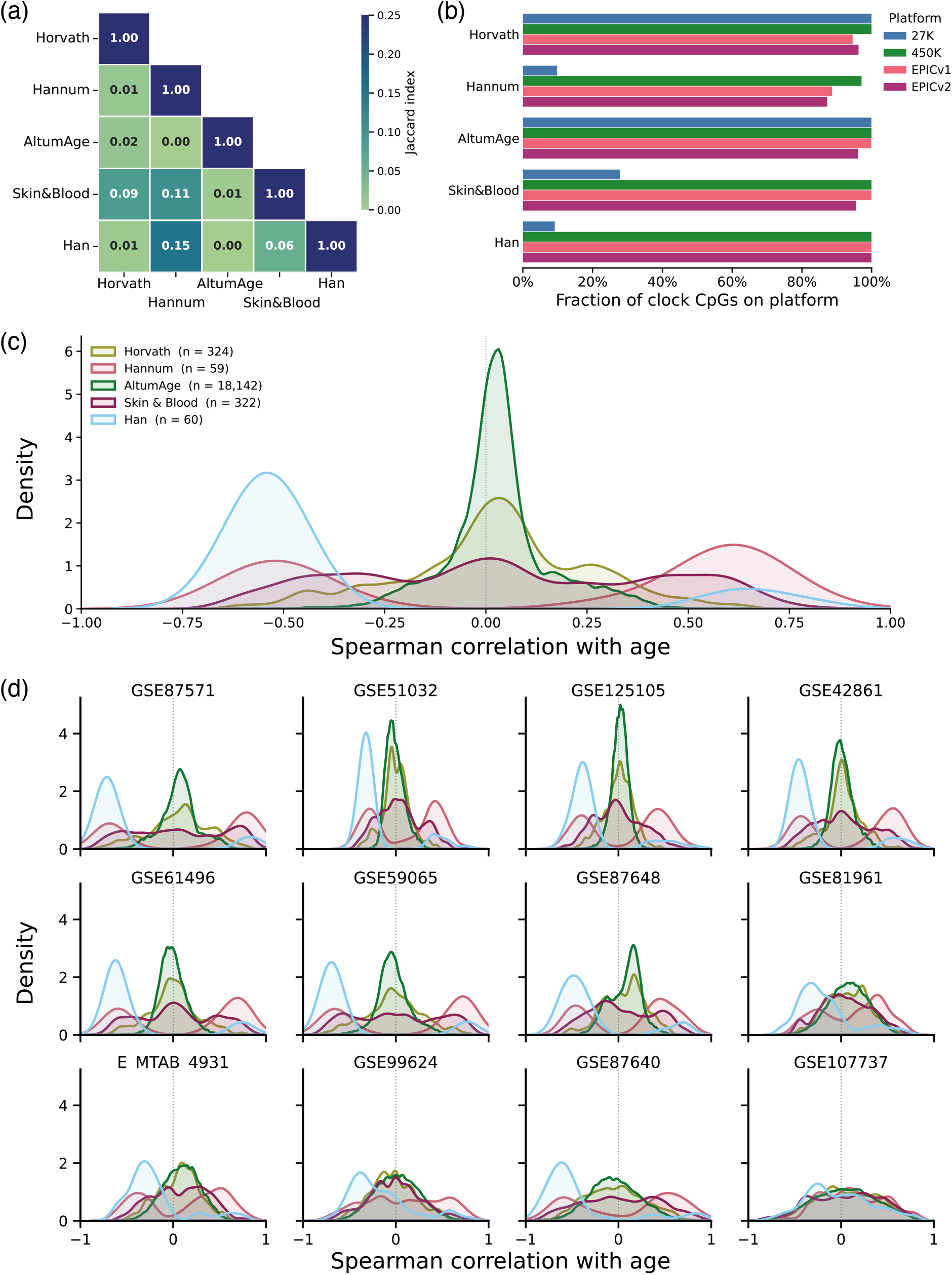
Limitations of existing DNAm clocks. (a) Pairwise CpG overlap across widely used clocks quantified by the Jaccard index. Most clock pairs share few or no CpGs (bright off-diagonal boxes). (b) Overlap between clock CpG sites and common Illumina methylation arrays. The overlap varies with Illumina platforms. (c) Spearman correlation values (*ρ*) with respect to chronological age for clock CpG sites using 12 datasets. The peaks at *ρ* = 0 show that many clock CpGs do not correlate with chronological age. (d) Spearman correlations with chronological age for individual datasets. There are variations between datasets, but many clock sites do not correlate with age.

A second observation is that clocks translate poorly across array platforms. The platforms evolve quickly, and each new generation drops some of the probes used by earlier clocks. To quantify this loss, we track the fraction of clock-associated CpGs that remains available from Illumina 27K through to 930K arrays (EPIC v2) (see Fig. 2b). The Hannum and Horvath clocks lose the most coverage, consistent with previous comparisons^51^. Only the Han clock retains full coverage in EPIC v2. The others retain enough probes to function, but their predictive power diminishes with platform generation^52^.

A third issue is that many individual clock CpGs show little correlation with chronological age. To show this, we compute the Spearman correlation *ρ* between *β* value and chronological age for every clock CpG, combining all 12 datasets profiled on 450K arrays (Fig. 2c). The distributions show a small fraction of CpGs with strong age associations (|*ρ*| *>* 0.6), but most are unaffected (*ρ* ≈ 0), particularly for Horvath, AltumAge, and, to some extent, Skin & Blood clocks. Hannum and Han clocks, by contrast, contain mostly sites that correlate strongly with age.

One explanation of this finding is tissue specificity. Hannum and Han clocks are trained on blood, so their CpGs are expected to correlate with age in our data sets, while Horvath and AltumAge clocks are fitted to data across many tissues and may include sites that are not relevant in blood. Skin & Blood clock falls between the two extremes, likely because it is trained on a narrower set of related tissues. This pattern holds across individual datasets (Fig. 2d), too, suggesting that it is not an artifact of any one cohort. However, tissue specificity is likely not the complete answer. The two blood-trained clocks still share only 15% of their CpGs, so even within a single tissue, very Different CpG collections can yield strong age predictions. Why so many low-correlation CpGs end up in the clocks at all remains an open question.

To sum up, existing clocks rely on largely non-overlapping CpGs, contain many sites with weak age signal, and do not transfer cleanly across array platforms. Together, these results highlight a few unresolved questions in the field.

### Networks of co-varying CpG sites are scale-free and modular

To investigate whether CpG sites organize into coordinated groups, we construct co-methylation networks from DNAm data using the 12 datasets. Because DNAm arrays contain thousands of CpGs, constructing a full co-methylation network would be computationally impractical. We therefore restrict the analysis to CpGs that change with age. To this end, we compute the absolute Spearman correlation with age for, and rank them as shown in Fig. 3a. Next, we retain CpGs with |*ρ*| ≥ 0.35 (dashed line in Fig. 3a), yielding approximately 7,000 sites for network construction (see details in *Materials and methods*). Although this threshold is arbitrary, varying it does not qualitatively alter the results (S7 Fig for results with |*ρ* | ≥ 0.3 and |*ρ* | ≥ 0.4).

**FIG. 3.**
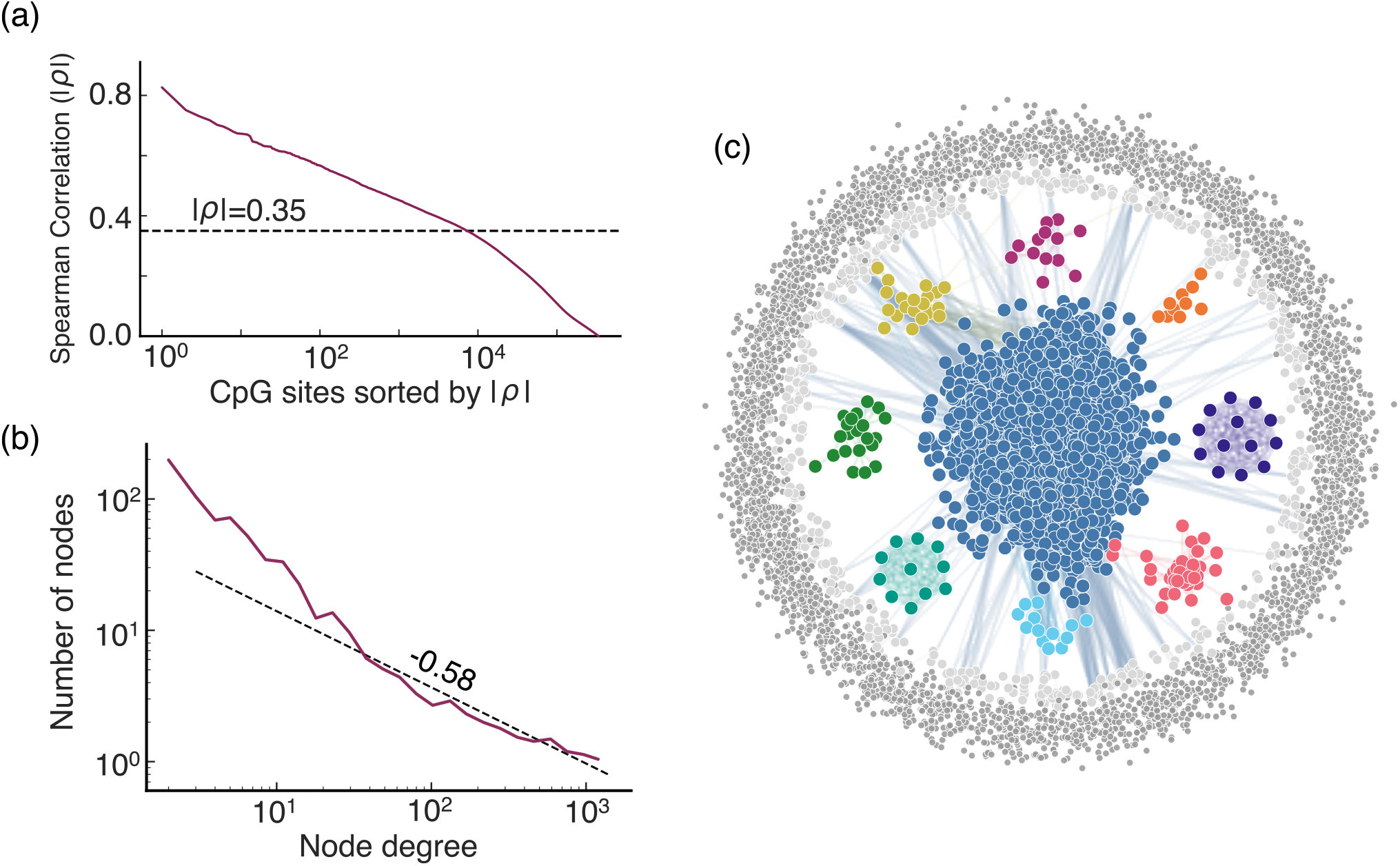
Node filtering and network clustering for the union of the 12 methylation datasets. (a) Ranked CpG sites according to Spearman correlation with age (*ρ*). We retain nodes with |*ρ* |*≥* 0.35. (b) Number of nodes as a function of node degree. The distribution is heavy-tailed and approximately follows a power law with slope 0.58 (dashed line). (c) Core network after filtering and pruning with color-coded modules extracted with Infomap.

The co-methylation network has CpGs as nodes, where two nodes are connected by a weighted link determined by the correlation between their respective *β* values. Since the correlation is non-zero in any practical setting, the resulting network is fully connected (with ∼ 2.5 ×10^7^ links). To extract a meaningful structure from the fully connected network, we therefore pruned edges using a threshold *τ* (see *Materials and methods*). This step removes insignificant links but also isolates a large number of nodes as “singletons” (≈ 62% with *τ* = 0.70). A separation of this kind, a strong core component and isolated nodes, is common in correlation-based biological networks, such as gene co-expression networks^37^. In addition, we find that the core contains a few well-connected nodes with orders-of-magnitude more links than other nodes (i.e., high node degrees). By plotting the nodes’ degree distribution, we find that it decays as a power law (Fig. 3b). Such a broad distribution is the hallmark of a heterogeneous scale-free network.

Next, we cluster the network. We found a pronounced modular organization that contains multiple well-defined communities with strongly correlated CpG sites (Fig. 3c), with one large module (blue) containing roughly 20% of the nodes and eight smaller modules with 10^1^–10^2^ nodes. The remaining nodes are either in small modules (∼ 10^3^ nodes) or in a sea of singletons (∼ 4 × 10^3^ nodes).

This modular structure is not specific to the merger of 12 datasets. Repeating the analysis on each cohort individually and on three independent EPIC v1 datasets yields networks with qualitatively the same topology: a dominant central module, several smaller modules, and a periphery of weakly connected and isolated sites (S8 Fig). Although the total network size varies between these datasets, a portion of age-associated CpGs do co-vary tightly enough to form well-defined modules, so coordinated methylation change is a real feature of the data, not an artifact of any one cohort.

### Clock CpGs aggregate in a few modules and spread across the singleton background

The modular organization of the co-methylation network suggests a possible explanation for the limited overlap between clocks: Different clocks might use the same co-methylation modules but pick Different CpG representatives from them. To test this hypothesis, we overlay the CpG sites of the five established clocks onto the co-methylation network and count how the CpGs of each clock distribute across modules, small modules, and singletons (Fig. 4a and S1 Table).

**FIG. 4.**
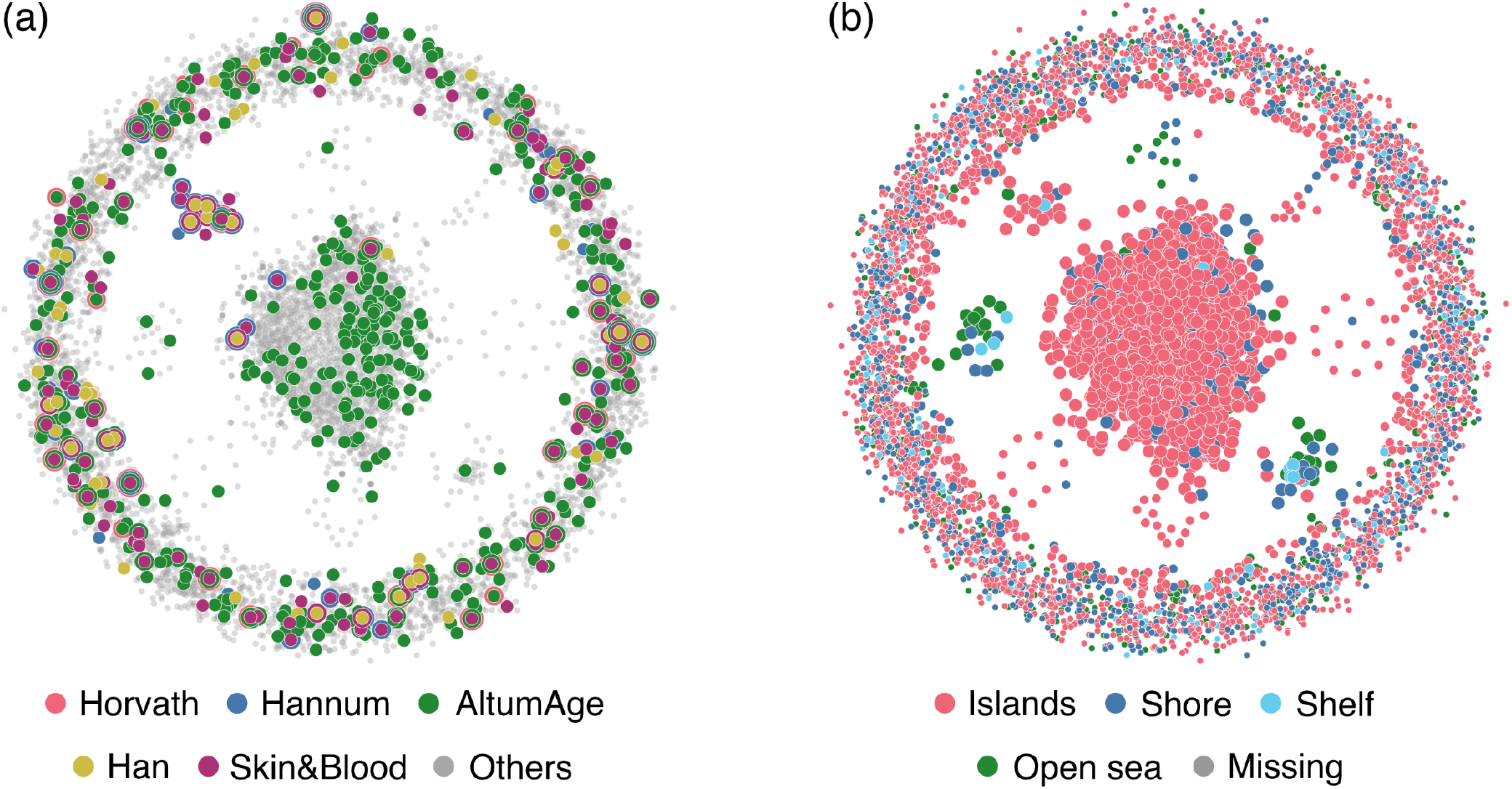
Co-methylation network with biological annotations. Nodes are age-associated CpGs and are colored depending on their additional biological information. (a) CpGs used by the five established clocks (color-coded), overlay on the network. Clock CpGs cluster on a few modules and scatter through the singleton periphery. (b) The same network coloured by CpG-island context. Each module is dominated by one context.

Most modules contain no clock CpGs at all. Most modules with more than 10 CpGs are empty across all five clocks. The exceptions are modules M1 and M4 (Fig. 3c, dark blue and yellow, respectively). M1 is the largest module with around 2, 000 CpGs, and it hosts CpGs from all clocks. The other module, M4 (with 20 CpGs), is also sampled by all five clocks. Note that the established clocks trained on blood samples and with the highest Jaccard overlap (Hannum and Han, Fig. 2a), draw most heavily from M4 (14.5% and 8.3% of their CpGs, respectively). The remaining large modules are not sampled.

The rest of the clock CpGs are in small modules with fewer than 10 CpGs, and the singleton background, where the singletons account for most of the clock CpGs: 89.4% of Horvath, 74.8% of Skin & Blood, 66.7% of Han, 65.5% of Hannum, and 59.0% of AltumAge. Small modules hold yet another 6%–22%. Every clock is therefore dominated by CpGs do not co-vary strongly with other CpGs, and share a minor contribution from specific modules.

### Network modules associate with CpG density

To test whether the modules align with known genomic features, we annotate each CpG site by its island context: island, shore or shelf flanking an island, or “open sea” away from islands (Fig. 4b). These categories track how densely CpGs are distributed along the DNA. We find that modules are strongly enriched for CpG islands. Of the 9 modules with ≥10 CpGs, five are dominated by island sites (S2 Table). M2 (Fig. 3, red) and M3 (green) are open-sea-dominated, with M2 also enriched for shore sites, and M6 (lilac) mixes shore and open sea.

We find the same pattern across the whole network. Large modules are the most island-dominated and small modules less so (71.7% islands). Singletons are the most mixed, with roughly half belonging to islands (54.9%), a quarter to shores (24.0%), and 16.1% to open sea (S2 Table). In other words, CpGs in large modules are more likely to be associated with (dense) CpG islands.

This analysis shows that the modular structure of the co-methylation network is not arbitrary. Each module tends to aggregate its CpGs from one or two genomic contexts, and the contrasts between modules separate islands from open-sea and shore regions.

### Modules capture biological functions

To assess whether the modules relate to distinct biological processes, we perform Gene Ontology (GO) enrichment analyses for each module (Fig. 5). For each module we keep the significantly enriched biological-process terms (nominal *p <* 0.05) and collapse related terms into broader parent categories. Across the four largest modules (M1– M4, each with more than 20 CpGs), these enriched terms do not overlap, and the separation holds even at the level of parent categories.

**FIG. 5.**
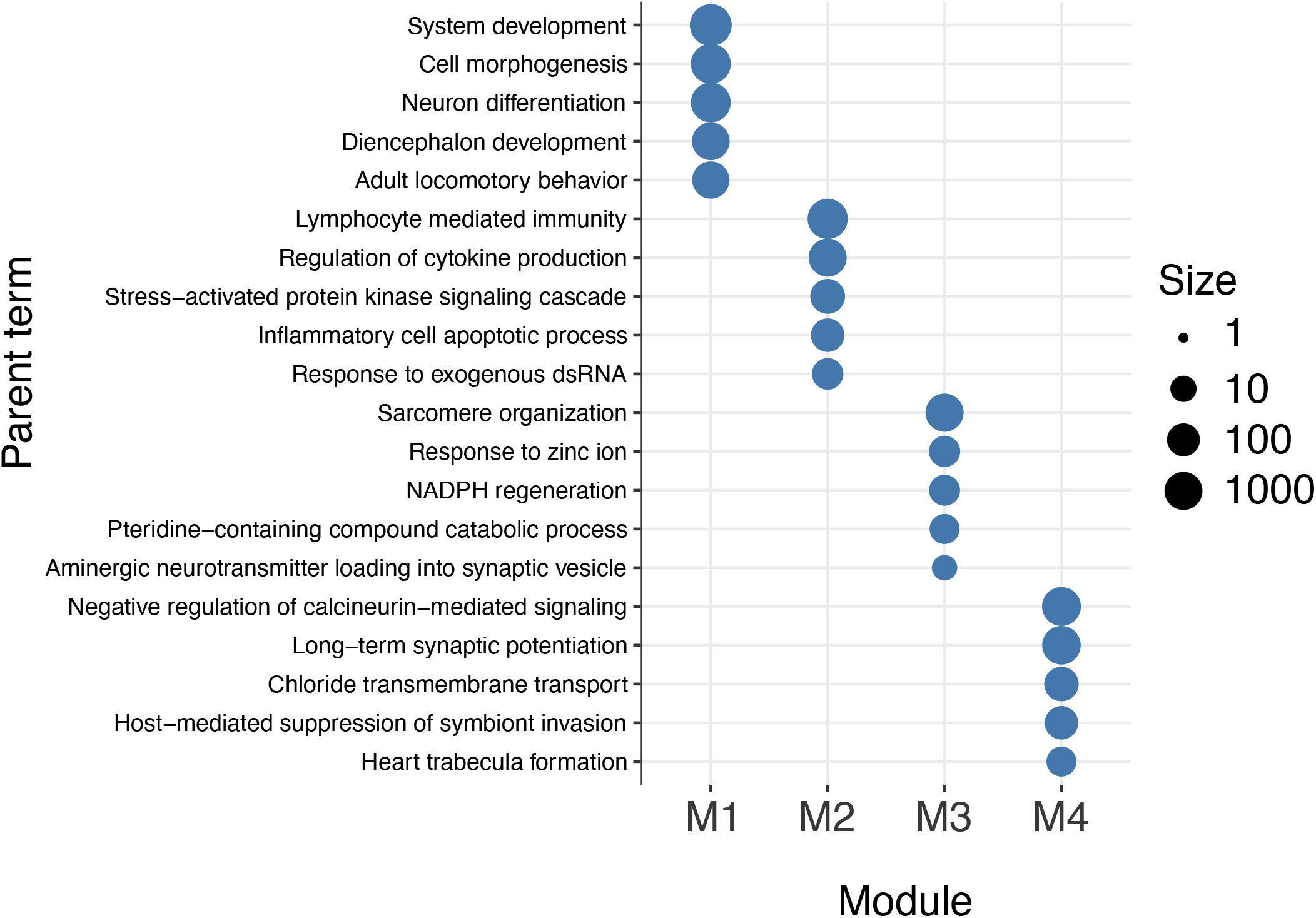
Gene Ontology enrichment across network modules. Each column is one of the four modules with more than 20 CpG sites (M1–M4), and each row is a parent GO term. We show the 5 most significant terms for each cluster. Dot size shows how many GO terms are collapsed into that parent term. The four modules are enriched with largely non-overlapping functions.

The modules differ in the kinds of processes they enrich for. Showing the five most enriched parent terms per module (Fig. 5), M1, the largest, spans broad neuronal and developmental themes, while M2, the second largest, enriches for immune and signaling processes; the smaller modules pick out narrower, more specific functions. None of these functional identities is shared between modules, reinforcing that the network separates age-associated methylation into distinct biological programs.

These results are exploratory. The enriched terms are significant at a nominal level but do not survive the stricter multiple-testing correction. Also note that GO enrichment on methylation data is indirect, since it uses CpG annotations as a proxy for gene function rather than measured expression^45;53^. Even so, the modules consistently enrich for separate functions, suggesting that age-associated methylation changes are spread across several biological processes rather than concentrated in one.

### A clock with one CpG per module matches established clocks

In the previous sections, we applied a network approach to understand genome-wide co-methylation. That analysis reveals a modular structure, which suggests a systematic way to select non-redundant sites for a new age predictor: a network clock. The idea is that a single CpG can represent the methylation state of its host module. We test it by randomly drawing *N* CpGs in two ways: either from anywhere in the network or at most once per module. Next, we use the methylation values for these sites to make a clock using Ridge regression.

Both strategies perform similarly well and surpass all reference clocks in the test data once *N* ≳ 200 (Fig. 6a). For comparison, the Hannum (blood trained) and Horvath (multiple tissues) clocks have 71 and 353 sites, respectively. One-per-module sampling consistently has a higher *R*^2^ value than random sampling for all *N*, though the difference is small, especially for large *N* . In this regime, both strategies sample the singletons, and the one-siteper-module constraint becomes irrelevant. However, for small *N*, this constraint affects site selection (compared to the uniform sampling) and is also the regime where we observe the largest differences between the two strategies. This advantage suggests the modules carry more signal than the singleton background. We investigate this further by selecting a single CpG per module, ordered by decreasing module size. This gives a clock that performs better than the one-per-module strategy (S10 Fig). Therefore, prioritizing the largest modules improves the clock and confirms that the larger modules drive accuracy.

**FIG. 6.**
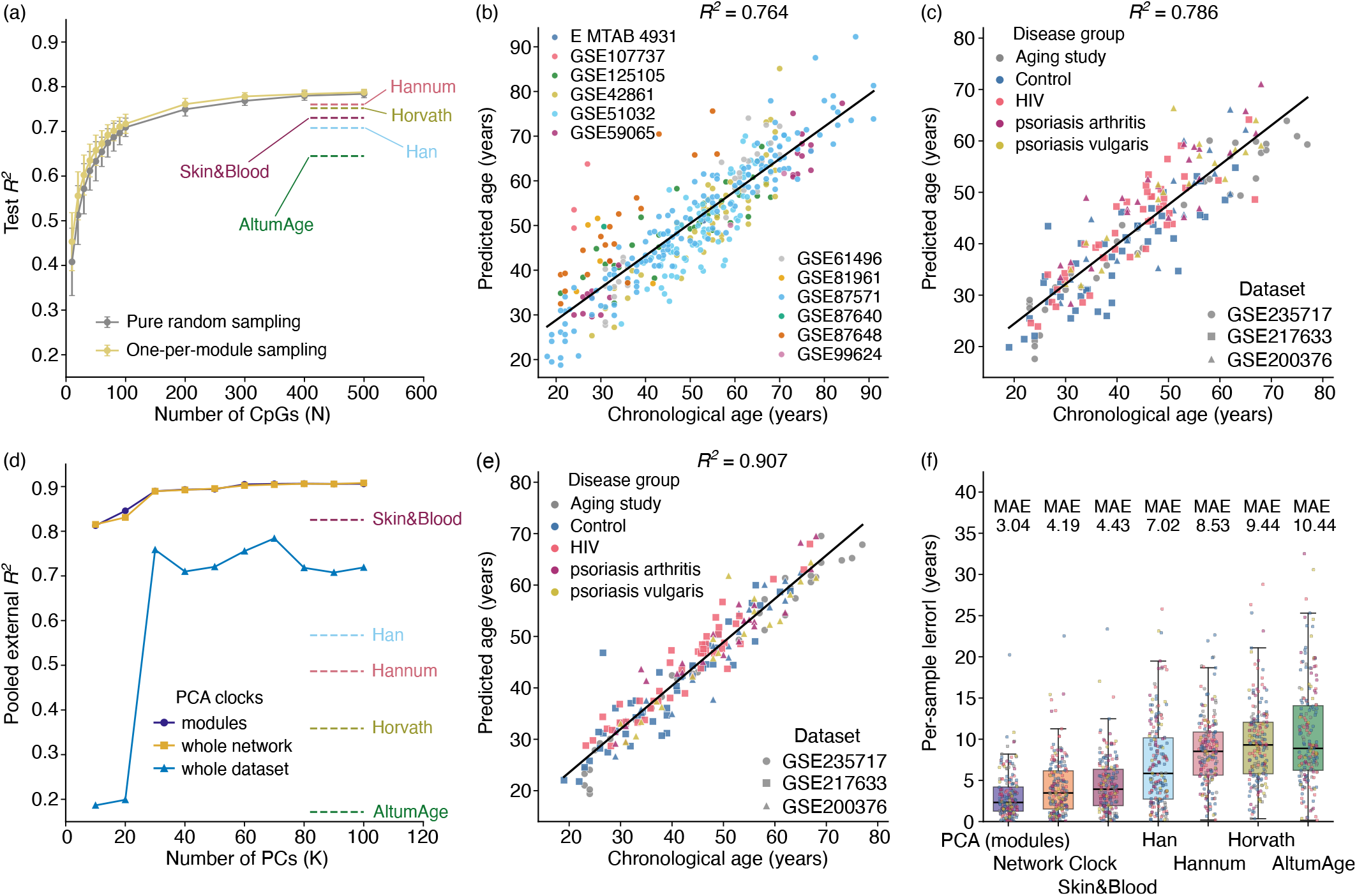
Network-based DNAm clocks match and surpass established clocks. (a) Test *R*^2^ versus number of CpGs *N*, for two network clock variants (random sampling and one-per-module sampling). The error bars indicate one standard deviation across 100 draws, and the solid lines are guides to the eye. The dashed horizontal lines show the reference-clock baselines. (b) Predicted versus chronological age on the test set for one network clock (*N* = 100, *R*^2^ ≈ 0.77). Coloured indicate dataset origin. (c) The best performing network clock applied to unseen validation sets (*N* = 500), *R*^2^ ≈ 0.79. Colors indicate disease group and dataset. This shows that network clocks transfer from 450K to EPIC v1 and applies across disease groups. (d) Test *R*^2^ against the number of principal components *K*, for three PCA clocks built on Different feature sets: the modules (dark blue), all age-filtered CpGs (yellow), and the full array (blue). (e) The best performing PCA clock applied to unseen validation sets (*K* = 60), *R*^2^ ≈ 0.90, as in (c). (f) Per-sample absolute error on the unseen validation sets [(c) & (d)] our best PCA and network clocks, and the five reference clocks. The module-based clocks give the lowest errors.

We also find that within a module, no particular site yields a better clock. For example, there is no difference between choosing the most central CpG by PageRank^54^ or one at random (S10 Fig). We conclude that CpGs within a module are exchangeable and a module contributes as a unit.

Next, we examine the predictions of one specific clock realization on the test data (one-per-module, 100 CpGs, *R*^2^ ≈ 0.77; Fig. 6b). We find a clear relationship between chronological age in each sample and its predicted age (albeit with a few outliers). We color-code the samples belonging to the same dataset. We note that a few of them are consistently above or below the regression line, likely due to cohort-specific biases, for example, a narrow age range or potential batch effects that could not be adjusted for.

To test the clock beyond the training data, we apply the best one-per-module clock (500 CpGs, *R*^2^ ≈ 0.83) to the held-out validation set with 98 samples from Different array platforms. On this data set, the network clock performs well. In addition, we included 89 samples with patients suffering from Different health conditions, such as HIV, Psoriasis Vulgaris, and Psoriatic Arthritis (Fig. 6c). The network clock predicts age about as well on this unseen data, both healthy and sick, as on the test samples. Its per-sample absolute error matches Skin & Blood clock but is lower than those of Horvath, Hannum, AltumAge, and Han clocks (Fig. 6f, second bar). Apart from being a good predictor of age, this result shows that the network clock is valid across platforms (here, 450K and EPIC v1).

### A module-level PCA clock outperforms reference clocks

A drawback of the network clock is that each realization randomly selects sites from Different modules, leading to variation between clocks, and we cannot tell which one generalizes best to the validation set. To mitigate this, we make a new clock following these two steps. First, we coarse-grain the data and represent each module by a single methylation value (the module median; see *Building a module-level PCA clock*, Methods). Second, we compute the principal components (PCs) of the coarse-grained dataset to build a new age predictor, the PCA clock, using a linear combination of the PCs (Eq. 5).

This module-based PCA clock outperforms all five reference clocks (Fig. 6d), including the network clock. We find that the *R*^2^ value increases with the number of PCs *K* on the held-out validation sets (for both healthy and sick) until it saturates at *K* ≥60, where *R*^2^ = 0.907 (dark blue line). Using the *K* = 60 clock, we show the explicit prediction for each sample (Fig. 6e) and the per-sample error (Fig. 6f). The PCA clock has the lowest median error among all tested clocks (Fig. 6f).

Next, we compare the module-based PCA clock with two reference cases. First, we compare it against a PCA clock built without the module coarse-graining step. That is, using the complete age-filtered dataset with ∼ 7, 000 CpGs (Fig. 6d, yellow line). We find that the module-based PCA clock is a few percent better for *K ≲* 30. But if more PCs are included, they reach similar performance. As a second reference case, we compare the module-based clock against one built on the full training array (all sites that pass normalization and filtering, ∼ 320, 000). The full-array clock improves sharply from *K >* 20 but has a much lower *R*^2^ value (Fig. 6d, blue line). It is also worse than Skin & Blood. This illustrates that including the whole array is not helpful, likely because adding too much noise.

Overall, age filtering is essential for good age prediction, and including the modules slightly improves performance for small *K*. To make the clock accessible and easy to use, we provide the source code and pipeline in a public repository^55^.

## IV. DISCUSSION AND CONCLUSION

Our results shed new light on recurring themes observed in DNA methylation clocks: they predict age well, yet are difficult to interpret. For example, they share few CpGs, do not transfer consistently across array platforms, and rely on CpGs that often seem uncorrelated with age. To better understand these observations, we created a co-methylation network from DNAm data curated from several datasets. We find that the network consists of a few large modules surrounded by a sea of singletons, where each module contains co-varying CpGs. These modules represent coherent methylation units that provide a coarse-grained representation and are easier to interpret than single CpG sites.

Our results suggest that much of the age signal captured by clocks is organized at the level of modules rather than independent sites. By mapping five established clocks onto the network, we find that most clock-associated CpGs belong to a few large modules, while the remaining sites fall into the singleton background. This means that the clocks are more similar than a simple CpG-CpG overlap analysis would suggest. The modular structure also suggests a more systematic way of building clocks. When we construct clocks using only one representative site per module, they achieve performance comparable to established clocks, almost regardless of which specific site we select. Beyond performance, such module-based clocks also translate better across array platform updates because modules remain stable under changes in probe coverage, whereas individual CpG sites may disappear. Another consequence of the modular structure is that it constrains the space of possible clocks. Instead of selecting clock sites from hundreds of thousands of CpGs across an array (e.g., using penalized regression), one e”ectively selects representatives from a much smaller set of co-varying modules. From this perspective, many existing clocks may be viewed not as fundamentally different models but as members of an ensemble of closely related members. Their comparable performance suggests that the predictive power resides primarily in the modules rather than in the specific CpG sites chosen to represent them.

The modules also have biological significance. First, each module is associated with a single CpG density type along the DNA (island, shore, etc.) As the density correlates with gene regulation activity, CpG islands often lie in promoter regions and have the potential to control gene expression via methylation^53;56^. This suggests that modules collect sites with similar regulatory roles. Second, the modules are also associated with specific gene functions (Gene Ontology terms). This suggests that each module maps to its own set of functions, with little overlap between modules. Together, these two patterns indicate that the modules capture biologically coherent and separable components of age-associated change.

In conclusion, our results suggest that the CpGs used in DNA methylation clocks are better understood through the structure of the co-methylation network than as independent predictors chosen in isolation. This modular organization explains how clocks with little CpG overlap can achieve similar predictive performance and why individual clock sites can be exchanged without substantially losing accuracy. The network perspective therefore shifts attention from individual CpG sites to the modules that organize them. By treating modules as the fundamental units of the aging signal captured with DNA methylation, we provide a framework that connects clock construction, interpretation, and biological context. This perspective may help explain why DNA methylation clocks remain robust despite their differences while providing a more systematic basis for developing the next generation of aging biomarkers.

## Data Availability

All code underlying this study is openly available on GitHub. The data preprocessing and network clock construction code is available at^57^. The DNA methylation datasets analysed in this study are publicly available (see citations in the text).

Additionally, we provide a GitHub repository with a ready to use pipeline for the PCA clock^55^.

## Acknowledgments

We thank Mattias Landfors for providing valuable feedback and support with the datasets. We also thank Rubén Bernardo Madrid for enriching discussions and ideas. S.-G.Y is supported by the Basic Science Research Program through the National Research Foundation of Korea (NRF), funded by the Ministry of Education (Grant No. RS-2025-02312897). ACM and LL acknowledge support from the Swedish Research Council (Grant No. 2021-04080). This study was supported by grants from The Swedish Childhood Cancer Foundation (SDE, SW), the Swedish Cancer Society (SDE), and the Cancer Research Foundation in Northern Sweden (SDE).

