## Supplementary Figures S1 to S10 and Tables S1 and S2 for "A Network Approach to DNA Methylation Clocks"

### V. SUPPORTING INFORMATION

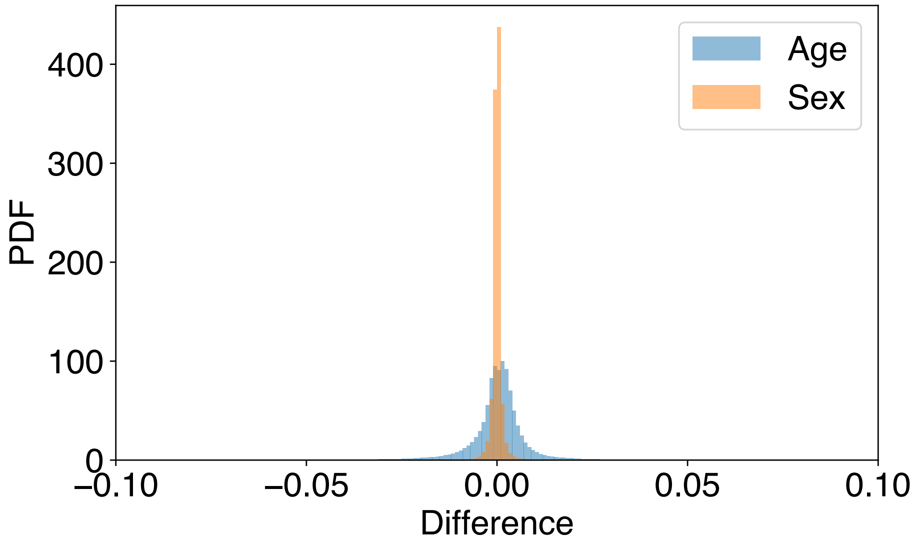


**S1 Fig. Distributions of difference in $\beta$ values.** We compare the beta values after ComBat with and without age and sex as biological covariates. With age as a biological covariate, the mean of the difference is $1.1 \times 10^{-4}$ and the standard deviation is $8.4 \times 10^{-3}$. For sex, the mean is around $-7.08 \times 10^{-7}$ and the standard deviation is $1.27 \times 10^{-3}$. For both, the mean is almost zero and the standard deviation is negligibly small. Since we filter out the sex chromosomes, the sharper distribution observed with sex as a biological covariate is expected.

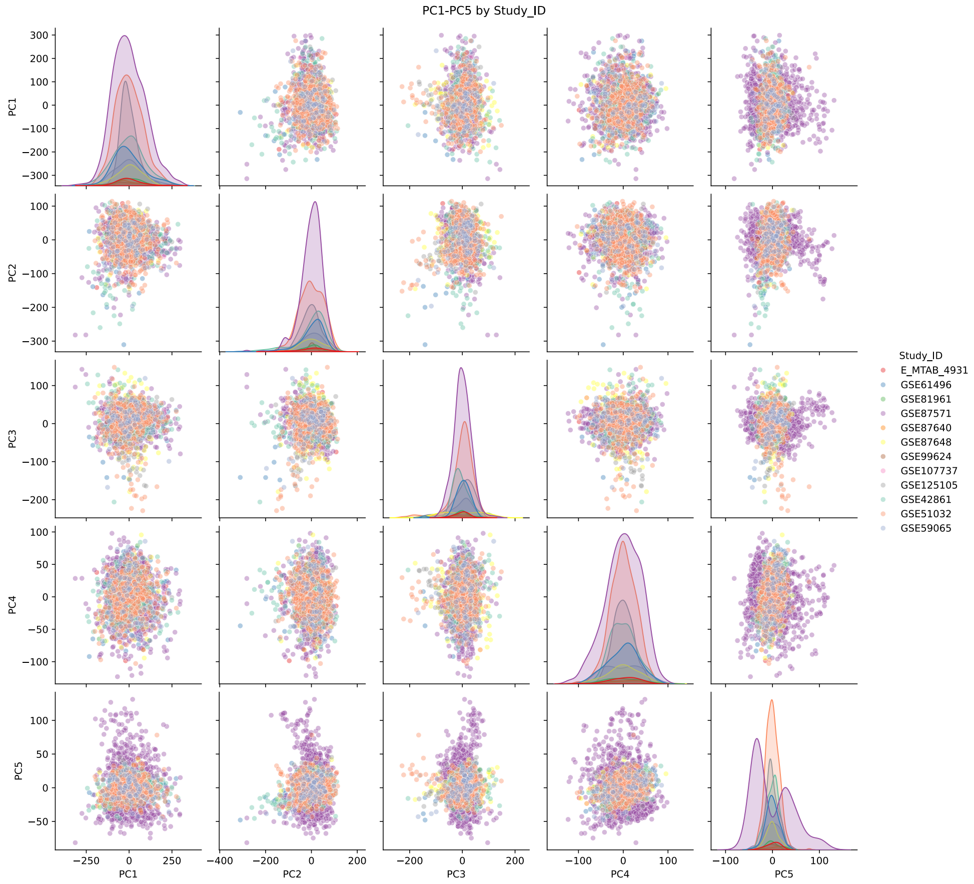

**S2 Fig. Looking at the PCs, we see no grouping by dataset after ComBat.** By PCA decomposition of the data, we check that ComBat does not introduce an artificial signal separating samples by their dataset.

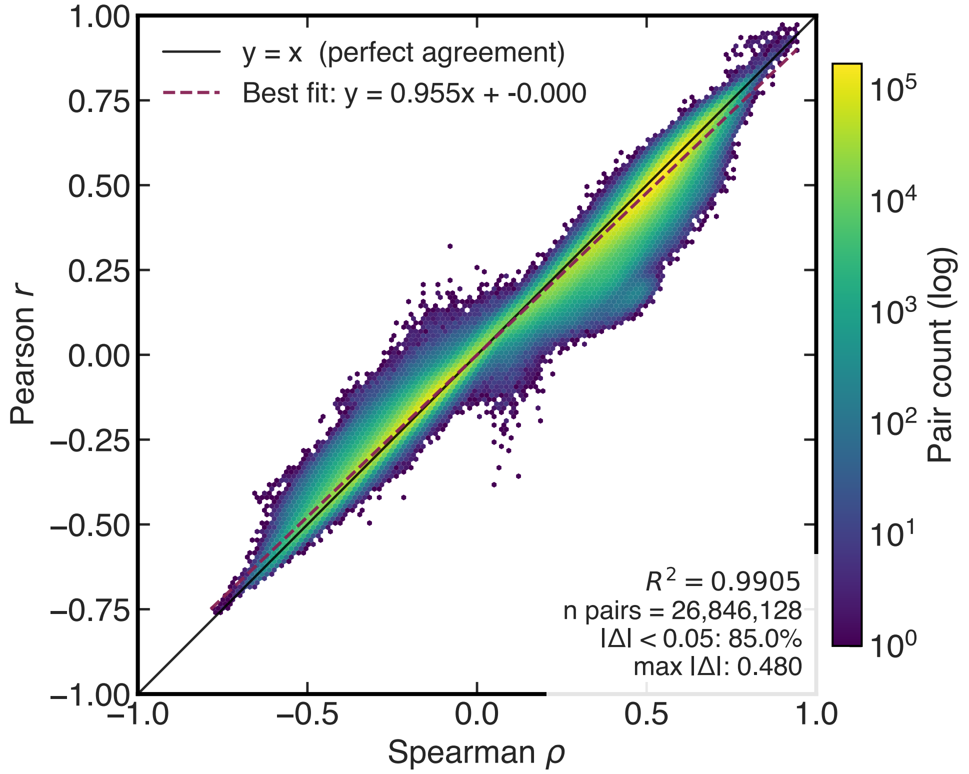


**S3 Fig. Pearson and Spearman age filtering agree.** CpG sites retained under $|\rho_{\text{age}}| \geq 0.35$ using Pearson versus Spearman correlation.

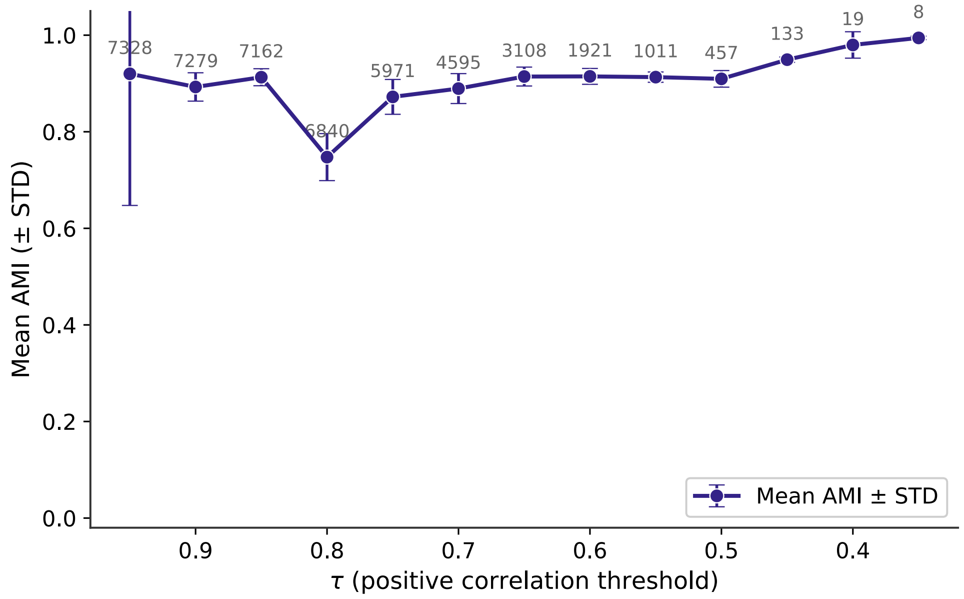


**S4 Fig. Threshold selection by partition stability.** Adjusted mutual information (AMI) between bootstrap partitions as a function of the correlation threshold $\tau$, with the number of modules and singletons overlaid.

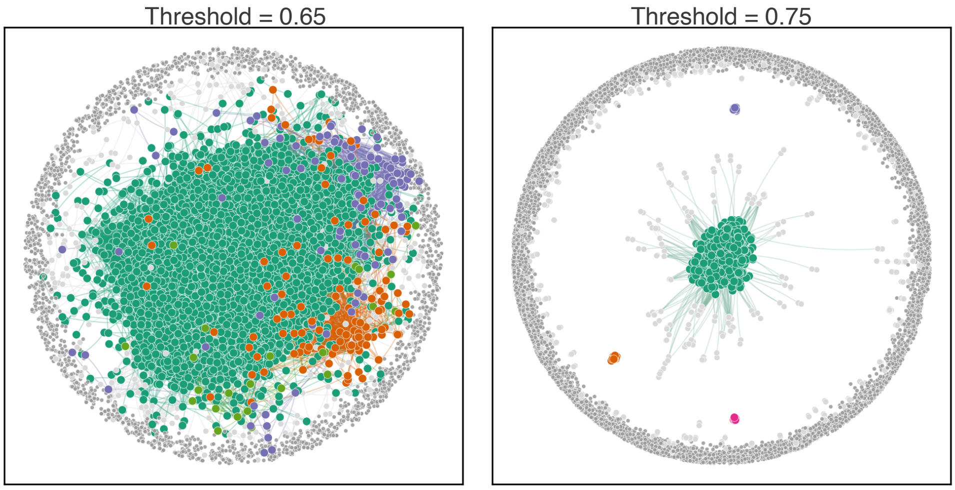


**S5 Fig. At $\tau$ ranging between 0.75 and 0.6 we get similar network structure.** When running links with $0.75 \geq \tau \geq 0.6$, we get big modules embedded in a sea of singletons with some smaller modules. The smaller the $\tau$, the bigger the biggest module is; increasing the threshold splits the big module into relevant submodules.

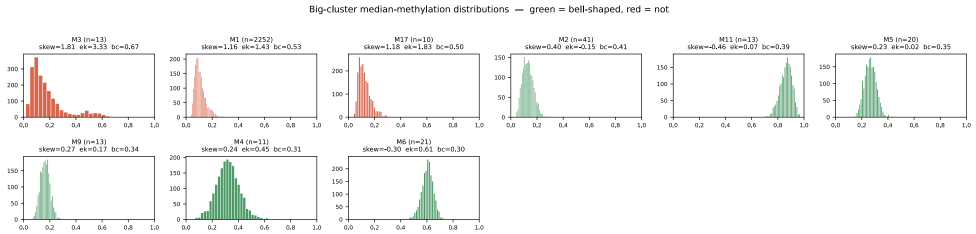


**S6 Fig. Modules are better represented by the median since outliers can skew the mean.** To get a representative methylation value from each module, we plot the distribution of the methylation of their CpGs. We see that they are bell-shaped, but with few CpGs far from the mean; therefore, the median is a more faithful representative value of the module.

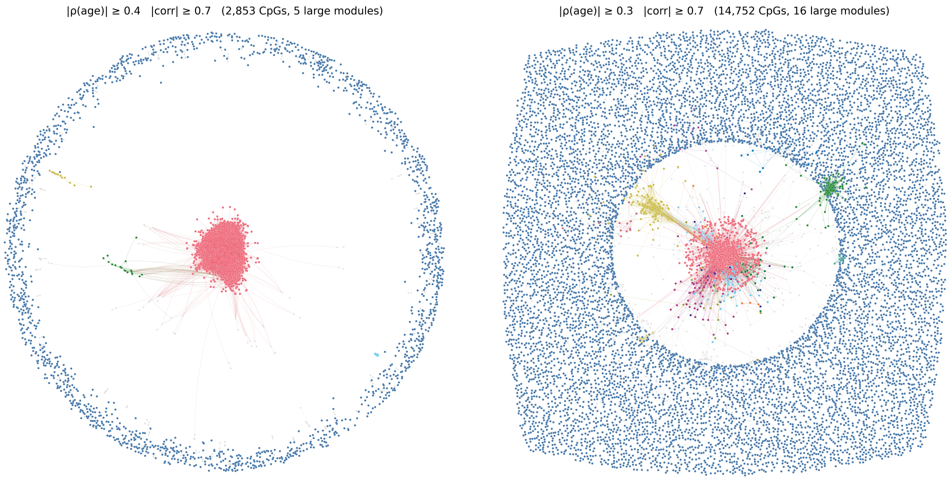


**S7 Fig. Varying $\rho$ does not alter the network structure.** We get the modules-surrounded-by-singletons structure, but the less restrictive the threshold, the more computationally demanding and the more CpGs that do not correlate with age.

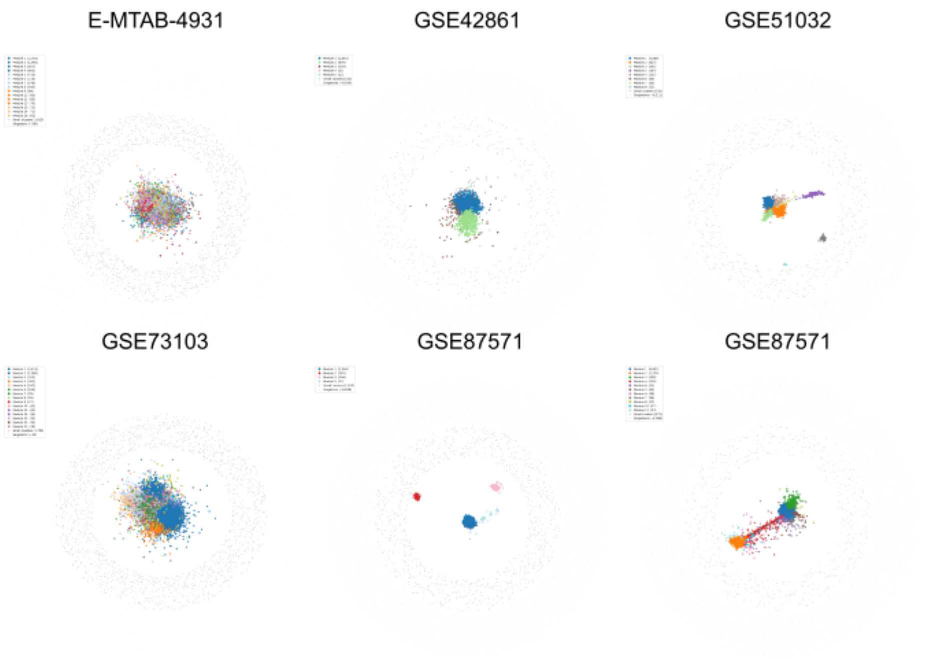


**S8 Fig. Different datasets give the same qualitative network structure.** We run the pipeline on other cross-sectional datasets of healthy individuals and obtain the same network structure.

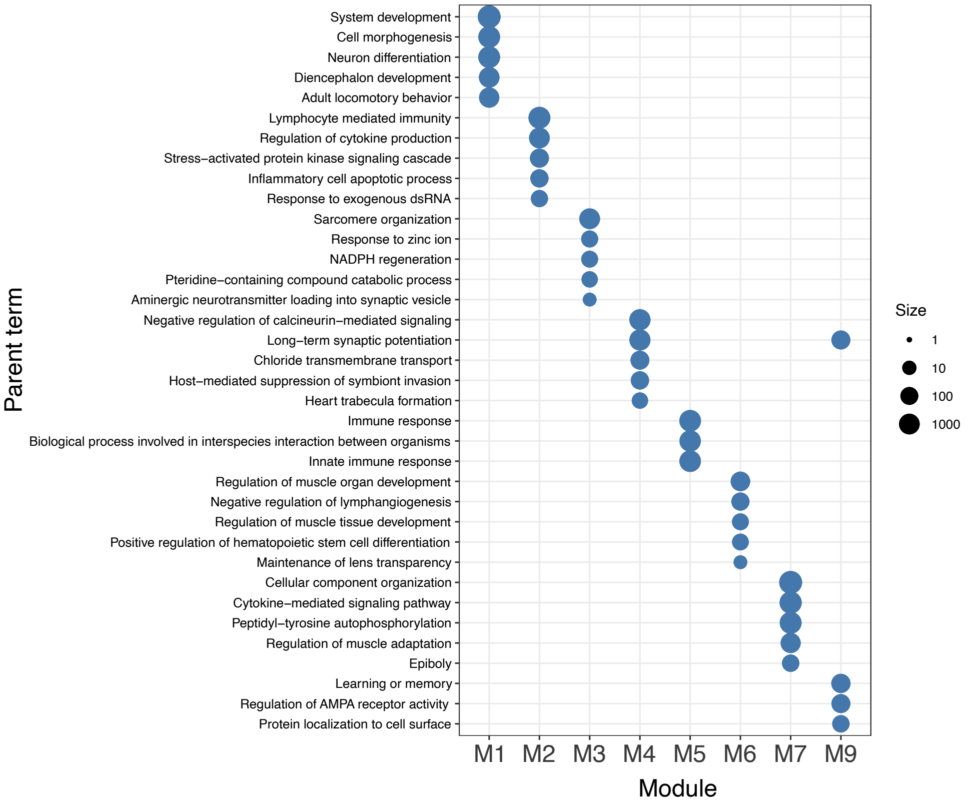


**S9 Fig. Gene Ontology enrichment across all network modules with 10 or more CpGs.** The modules enrich for non-overlapping functions, as seen for the larger set ($\geq$ 20 CpGs) in Fig. 5.

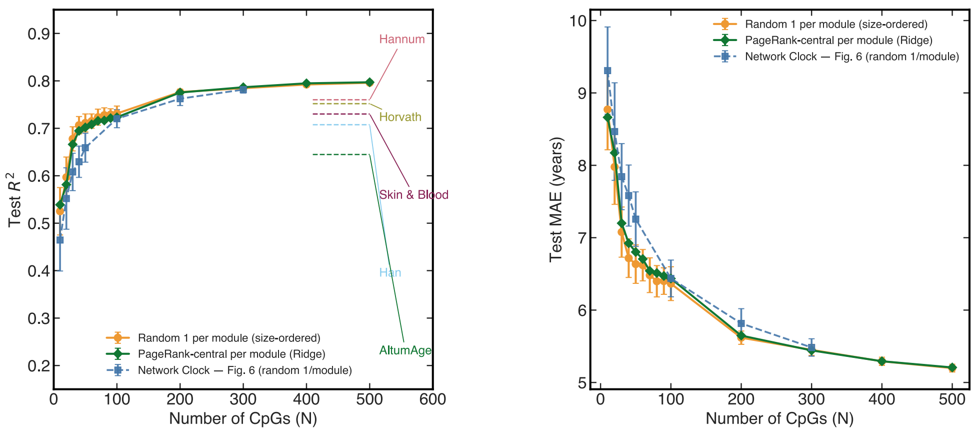


**S10 Fig. Modules carry more signal than singletons, and CpGs within a module are exchangeable.** Test $R^2$ versus the number of selected CpGs $N$ for clocks built in different ways. Random one-per-module sampling (one CpG drawn at random from each module) is compared with two deterministic constructions that add module representatives in order of decreasing module size before the age-ranked singletons, choosing each representative either at random or as the most central CpG of the module by PageRank. The size-ordered constructions, which prioritise the largest modules and use no sampling, perform slightly better than random one-per-module sampling, indicating that the modules carry more age signal than the abundant singleton sites. The two size-ordered constructions (random and PageRank CpGs) are nearly identical, showing that the choice of representative CpG within a module does not matter and that the CpGs in a module are interchangeable for age prediction.

| | Horvath (47) | Hannum (55) | AltumAge (461) | Skin&Blood (143) | Han (60) |
|---|---|---|---|---|---|
| M1 | 1 (2.1%) | 3 (5.5%) | 146 (31.7%) | 5 (3.5%) | 2 (3.3%) |
| M2 | 0 (0.0%) | 0 (0.0%) | 2 (0.4%) | 0 (0.0%) | 0 (0.0%) |
| M3 | 0 (0.0%) | 0 (0.0%) | 3 (0.7%) | 0 (0.0%) | 0 (0.0%) |
| M4 | 1 (2.1%) | 8 (14.5%) | 1 (0.2%) | 12 (8.4%) | 5 (8.3%) |
| M5 | 0 (0.0%) | 0 (0.0%) | 0 (0.0%) | 0 (0.0%) | 0 (0.0%) |
| M6 | 0 (0.0%) | 0 (0.0%) | 1 (0.2%) | 0 (0.0%) | 0 (0.0%) |
| M7 | 0 (0.0%) | 0 (0.0%) | 0 (0.0%) | 0 (0.0%) | 0 (0.0%) |
| M8 | 0 (0.0%) | 0 (0.0%) | 1 (0.2%) | 0 (0.0%) | 0 (0.0%) |
| M9 | 0 (0.0%) | 0 (0.0%) | 0 (0.0%) | 0 (0.0%) | 0 (0.0%) |
| Small modules | 3 (6.4%) | 8 (14.5%) | 35 (7.6%) | 19 (13.3%) | 13 (21.7%) |
| Singletons | 42 (89.4%) | 36 (65.5%) | 272 (59.0%) | 107 (74.8%) | 40 (66.7%) |

**S1 Table. Clock sites per module.** The number of clock sites in each large module (10 or more CpGs), with the small modules and singletons listed as separate groups.

| | Island | Shore | Shelf | OpenSea |
|---|---|---|---|---|
| M1 (2,252) | 2,042 (90.7%) | 166 (7.4%) | 5 (0.2%) | 39 (1.7%) |
| M2 (41) | 6 (14.6%) | 13 (31.7%) | 5 (12.2%) | 17 (41.5%) |
| M3 (21) | 0 (0.0%) | 5 (23.8%) | 3 (14.3%) | 13 (61.9%) |
| M4 (20) | 17 (85.0%) | 1 (5.0%) | 1 (5.0%) | 1 (5.0%) |
| M5 (13) | 12 (92.3%) | 1 (7.7%) | 0 (0.0%) | 0 (0.0%) |
| M6 (13) | 1 (7.7%) | 5 (38.5%) | 0 (0.0%) | 7 (53.8%) |
| M7 (13) | 13 (100.0%) | 0 (0.0%) | 0 (0.0%) | 0 (0.0%) |
| M8 (11) | 9 (81.8%) | 2 (18.2%) | 0 (0.0%) | 0 (0.0%) |
| M9 (10) | 8 (80.0%) | 2 (20.0%) | 0 (0.0%) | 0 (0.0%) |
| Small modules (583) | 418 (71.7%) | 86 (14.8%) | 6 (1.0%) | 73 (12.5%) |
| Singletons (4,351) | 2,387 (54.9%) | 1,045 (24.0%) | 218 (5.0%) | 701 (16.1%) |
| Network (vs 450K) (150,254) | 4,913 (3.3%) | 1,326 (1.2%) | 238 (0.5%) | 851 (0.5%) |

**S2 Table. CpG density context per module.** The CpG-density context in eachlarge module (10 or more CpGs), with the small modules and singletons listed as separate groups.
